# Gene functional-state covariance reveals modular structure in the *Arabidopsis thaliana* dispensable genome

**DOI:** 10.64898/2026.10.08.757691

**Authors:** Kevin A. Bird, Kehan Zhao, Daniel J. Kliebenstein, J. Grey Monroe

**Affiliations:** Department of Trait Diversity and Function, Royal Botanic Gardens, Kew; Department of Plant Sciences, University of California, Davis

## Abstract

Classical multilocus theory predicts that selection acting on combinations of loci can maintain non-random associations among allelic states, causing selection to act on multilocus genotypes rather than individual alleles. Across eukaryotic species, pangenome studies have documented thousands of genes that vary in functional status between individuals, yet this variation is analyzed as if each gene evolves independently. Here we test for non-random association within the dispensable pangenome across 1,012 natural genotypes of *Arabidopsis thaliana* to better understand the structure and evolution of eukaryotic pangenomes. Genome-wide pairwise association testing of loss-of-function alleles across genotypes identified 150,033 significantly covarying gene pairs among 5,679 unique genes after controlling for kinship and population structure. Most associations were between genes on different chromosomes, and covarying gene pairs showed greater co-expression and functional similarity than expected by chance. Network analysis revealed a non-random higher-order architecture comprising stable functional communities with distinct biological functions and structured connectivity among communities. The enriched functions and connectivity of communities revealed a tight coupling of modules involving plant immunity, environmental response, and specialized metabolism, highlighting how these systems integrate and evolve in response to dynamic ecological pressures. Taken together, these results show that the dispensable genome has a complex higher-order structure driven by functionally interacting genes, and that variation in gene content provides a population-scale resource for investigating the organization and evolution of biological systems.

## Introduction

An outstanding question in genetics is how the mechanistic architecture controlling traits shapes patterns of genetic variation in a species. Multilocus population genetic theory predicts that selection can generate non-random associations among combinations of physically unlinked loci, such that selection acts on multilocus genotypes rather than individual alleles (Lewontin and Kojima, 1960; Neher and Shraiman, 2009). Ecological geneticists independently arrived at a similar conclusion from field data, noting that the way locally adapted populations differed simultaneously in morphological and physiological traits suggested that selection acts on suites of characters regulated by systems of genes, rather than individual genes (Clausen and Hiesey, 1960). Adaptation, in this view, is a property of coherent genotypes rather than of independently favorable alleles. Empirical studies across model organisms have lent support to these predictions, identifying strong evidence that mechanistic interactions and associated epistatic variance structure complex trait variation at the population level (Bloom et al. 2013; Clegg et al. 1972; Hartl, 2014; Huang et al. 2012; Kerwin et al. 2017; Rowe et al. 2008; Weinreich et al. 2013). However, this body of work has largely failed to penetrate studies of pangenomic variation in eukaryotes. Pangenome studies have documented thousands of genes present and functional in some individuals but absent or non-functional in others, yet this variation is typically catalogued and analyzed as though the evolutionary fate of each gene is independent of the rest.

Contrary to this gene-by-gene framing, the presence of substantial epistatic variance implies that genomes could contain modular assemblies of co-selected loci whose internal coherence is maintained by their functional interaction (Brandon, 1999; Dobzhansky, 1950). Indeed, well-characterized naturally variable pathways in plant biology illustrate this modular co-adaptation. In *Arabidopsis thaliana*, interactions between the flowering regulators FLC (flowering locus C) and FRI (Frigida) contribute to adaptive flowering behavior, while covariation among additional flowering-time loci suggests broader multilocus organization (Michaels et al. 2003; Shindo et al. 2006; Zan and Carlborg, 2019). In the glucosinolate pathway, epistatic interactions among GS-ELONG, GS-AOP, and GS-OH create biochemical dependencies among loci, producing non-random combinations of functional and non-functional alleles in natural populations (Brachi et al. 2015; Katz et al. 2021; Kliebenstein et al. 2001). In both cases, epistatic/mechanistic relationships among these genes results in the overrepresentation of specific functional/non-functional allelic combinations to facilitate local adaptation across environmental clines (Caicedo et al. 2004; Hansen et al. 2008; Kerwin et al. 2015,2017; Züst et al. 2012). Beyond individual pathways, crossing experiments in Arabidopsis have demonstrated that the nucleotide-binding leucine-rich repeat (NLR) gene family involved in plant immunity contributes to widespread deleterious epistasis, where combining incompatible alleles results in a cell death auto-immune response due to the mechanistic connections between these genes (Chae et al. 2014). While these examples illustrate that multilocus organization can arise from both functional interdependence among genes and from incompatibilities between them, they are also drawn from specific, well-characterized pathways. Whether comparable organization is a general feature of genome-wide variation in gene content remains unknown.

Insights into the role of genetic interactions, their higher-order structure, and their effect on genetic variation have begun to emerge from recent work on bacterial pangenomes. In both *Pseudomonas* sp. and *E. coli,* a substantial portion of the accessory genome shows correlated gene presence/absence variation (PAV) structured by underlying biological networks and genetic interactions into functional modules (Whelan et al. 2021; Beavan et al. 2024). Simulation studies seem to confirm that these module assemblages require epistatic interactions to persist, and modules are only transiently present under neutral processes alone (Castro and Brown, 2022). Together, these studies suggest that functional interactions among genes can impose higher-order structure on pangenome variation, such that the evolutionary fate of one accessory gene depends in part on the presence or absence of other genes as the populations respond to dynamic environmental challenges (McInerney, 2023). However, bacterial pangenome variation is primarily driven by horizontal gene transfer, and it remains debated if eukaryotic gene content variation is primarily adaptive or largely neutral (Baumdicker et al. 2012; Bush et al. 2014). As yet, we lack a clear answer as to whether eukaryotic pangenomes, where gene content varies through duplication, transposition, and deletion rather than lateral transfer (Golicz et al. 2020), show the same epistatic/modular co-evolutionary structure as bacterial pangenomes.

If functional interactions among genes shape their evolutionary trajectories, functional states should not be distributed independently across natural populations. Instead, genes participating in shared biological processes should exhibit correlated loss or retention, potentially extending across physically unlinked loci. A substantial challenge is how to distinguish between covariance attributable to population history or genomic structure from covariance that reflects functional relationships among genes, and whether pairwise relationships resolve into higher-order organization. A network framework provides a means to do this by shifting the focus from individual associations to the structure formed by many associations. Here we test whether the dispensable genome of *Arabidopsis thaliana* exhibits higher-order structure reflecting the functional interactions among genes, as predicted by multilocus theory. Using naturally segregating loss-of-function variants from 1,012 accessions as a proxy for gene functional state, we performed genome-wide pairwise association testing while controlling for kinship and population structure, and then asked whether the resulting associations were concentrated among functionally related genes and organized into higher-order network communities. We find extensive covariance among predominantly unlinked genes, with associated genes showing greater functional and transcriptional similarity than expected by chance. The resulting network could be partitioned into distinct functional communities with structured connection, demonstrating that functional interactions among biological processes shape pangenomic variation.

## Methods

### LoF calling and LoF allele matrix

The detailed LoF calling pipeline is described in Zhao et al. (2026). Gene loss was identified from homozygous loss-of-function variants called from publicly available Whole-Genome Sequencing (WGS) data, including SNP variants and small indel calls for 1,135 Arabidopsis accessions from the 1001 Genomes Project (Alonso-Blanco et al.m 2016) and structural variants and indel calls for 1,301 accessions (Göktay et al. 2021). The VCF from the 1001 Genomes Project annotated using SnpEff was downloaded from 1001 Genomes Data Center (https://1001genomes.org/data/GMI-MPI/releases/v3.1/1001genomes_snpeff_v3.1/). The program snpEff (Cingolani et al. 2012) is widely used to annotate mutation effects from genotype variants. Variants annotated as “High” impact are considered as putative LoF variants. Those include “stop_lost”, “stop_gained”, “start_lost”, “frameshift”, “splice_donor_variants”, and “splice_acceptor_variants”. For structural variants, we mainly focus on deletions as they constitute the majority of the data. A deletion variant is considered as LoF if it induces a frameshift mutation predicted to disrupt the reading frame of the transcript or removes more than 10% of the protein-coding sequence of the affected transcript. It is also assumed that the protein product could still be functional with a change in the first and last 5% of the coding sequence as an alternative start/stop codon may rescue the disrupted transcript, and thus variants that only affect the first and last 5% of the coding sequence were therefore systematically removed (MacArthur et al. 2012). Lastly, to ensure true knockout of a gene and reduce false positives, only variants that disrupt all gene models for the gene of interest are kept. With the prediction of LoF alleles, a LoF allele matrix is generated to reflect the relationship between accessions and LoF alleles. Independent LoF alleles of the same gene were then collapsed into a single allele state. The collapsed LoF allele matrix represents a binary state of LoF or functional for each accession-gene combination, where LoF alleles are shown as “1” and functional alleles are shown as “0” in the LoF allele matrix.

### Genome-wide association of gene co-occurrence and avoidance

Genome-wide association testing based on pairwise gene LoF burden was performed using the EMMAX algorithm (Kang et al. 2010) implemented through the function cGWAS.emmax in the R package cpgen. The collapsed binary LoF allele matrix was used as the marker matrix, and for each association test, the LoF status of a focal gene from the same matrix was treated as the phenotype. A kinship matrix derived from SNP data from the 1001 Genomes dataset (https://1001genomes.org/data/GMI-MPI/releases/v3.1/SNP_matrix_imputed_hdf5/) was incorporated to account for population structure. The analysis produced both a p-value matrix and a beta-coefficient matrix, recording the significance and direction of each pairwise association. Positive beta values indicate gene co-occurrence, whereas negative beta values indicate gene avoidance.

### Data Filtering

To reduce false positives, association results were subjected to several filtering steps. First, p-values for each association were adjusted for multiple testing using the FDR method via the p.adjust() function, and associations with FDR-corrected p < 0.05 were retained. We filtered genes where the frequency of non-functional status was below 5% or above 95% to remove extremely rare variants. Gene positions were determined as the midpoint of the Araport11 gene model.

### Principal component analysis

Principal component analysis was performed on the collapsed binary LoF matrix using the prcomp function in R, with variables centred but not scaled. Two analyses were performed. The first included all accessions and was used to visualise the separation of the North American lineage from Eurasian accessions (Fig. 1D). The second was restricted to the 1,012 Eurasian accessions retained for association testing, and its accession scores were used in the community burden analyses described below (Fig. 6D).

**Figure 1.**
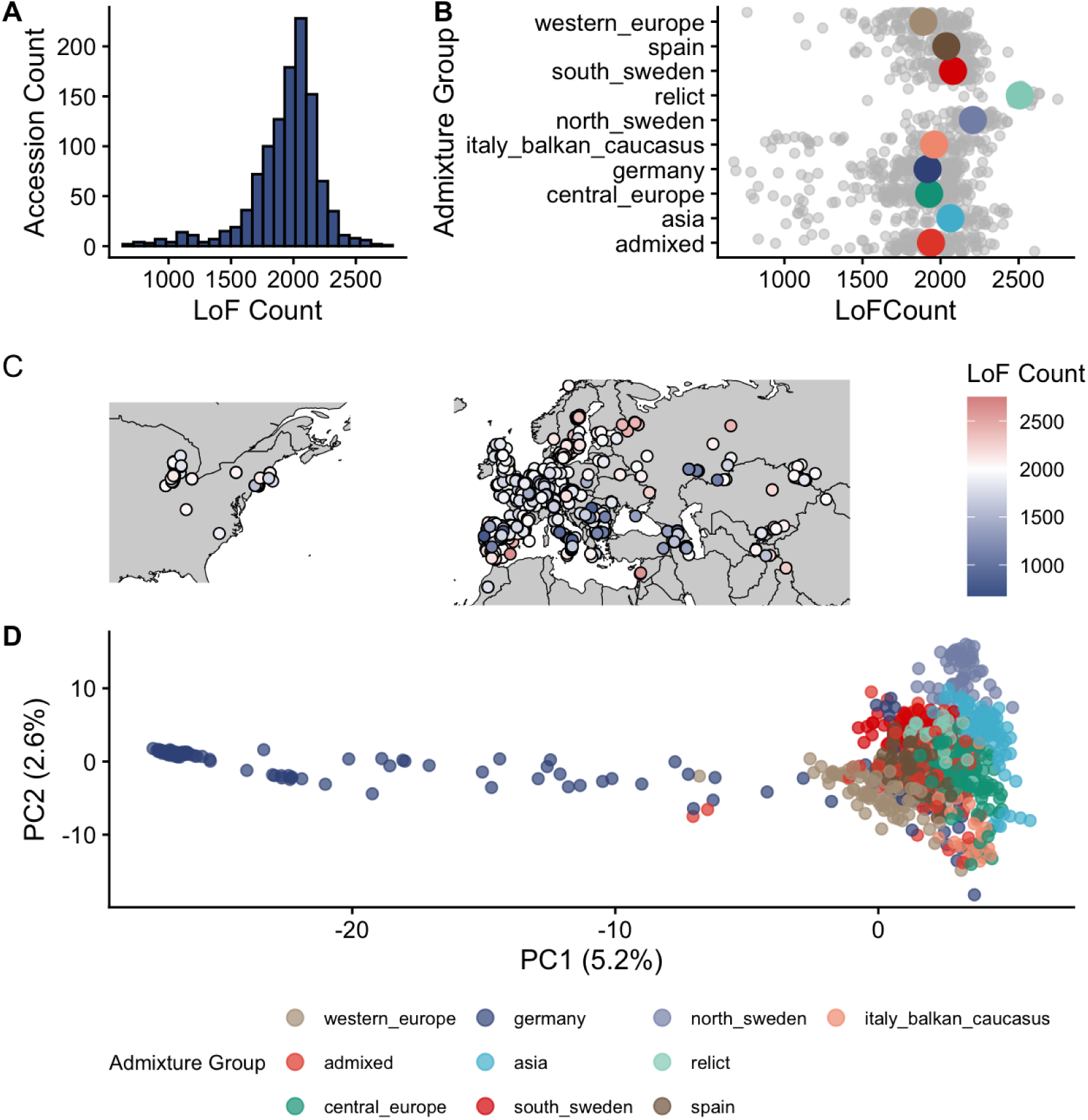
Functional state variation across global *Arabidopsis thaliana* accessions. A) The distribution of loss-of-function counts across genotypes. B) Variation in loss-of-function count across previously identified admixture groups. C) Geographic distribution of Loss-of-function counts across the population. D) Principal component analysis of loss-of-function profiles. Colors represent previously identified admixture groups.

### North American Accession Characterization

Accessions belonging to the North American HPG1 lineage were identified based on previous work by Exposito-Alonso et al. (2018). Genes fixed for the loss-of-function state within this lineage were defined as those with a LoF frequency of 1 across all HPG1 accessions. To test whether the gene losses fixed in this lineage were functionally unusual relative to standing variation elsewhere in the species, we compared the gene content of HPG1 accessions against that of Eurasian accessions in terms of selective constraint (Dn/Ds and neutrality index), expression breadth, and lethality of the corresponding knockout. North American accessions were excluded from all subsequent association testing.

### Pairwise coexpression analysis

To test for enriched coexpression among gene pairs with significantly functional-state covariation, we computed pairwise coexpression using the expression data from high-resolution developmental transcriptome map of Klepikova et al. (2016). For each gene pair coexpression was calculated as the Spearman rank correlation of their expression profiles across all treatments, developmental stages, and tissues in the atlas. We repeated this for each significant gene pair from the EMMAX analysis. We compared the observed mean coexpression for all significant associated gene pairs against a null distribution based on the mean coexpression of 100 permutated sets made by randomly shuffling gene pairs. This analysis was performed separately for positively and negatively associated gene pairs.

### Functional annotation overlap

Functional annotation overlap between significant gene pairs was assessed following the approach of Whelan et al. (2021). GO Biological Process annotations for all Arabidopsis genes were obtained from TAIR for Araport11, resulting in 3,703 unique biological process (BP) terms across 22,782 annotated genes. We first removed the highest level GO terms such as GO:0008150 (Biological Process). For each significantly associated gene pair with existing BP annotations, overlap was defined as the presence of at least one shared GO BP term between the two genes’ annotated terms. The observed proportion of overlapping pairs was compared against 100 permutations in which gene pairs were randomly shuffled, as the coexpression analysis permutation. The overlap analysis was implemented using the go_term_stats_perm_test.py script adapted from the Coinfinder pipeline (Whelan et al. 2021). This analysis was performed separately for positively and negatively associated gene pairs.

### Network construction

Significant positive associations from the EMMAX gene-by-gene analysis were used to construct an undirected weighted network, where nodes represent genes and edges represent significant co-occurrence relationships (FDR < 0.05, positive beta coefficient). Edge weights were set to the beta coefficient from the EMMAX mixed linear model. Only genes passing the joint allele frequency filter (both genes in a pair with LoF frequency between 0.05 and 0.95 in the European population) were included. The resulting network contained 5,673 nodes and 140,568 edges. Network construction and all downstream analyses were performed using the igraph package (v 2.2.2) in R (4.4.1).

### Network topology characterization

Network topology was characterized by computing transitivity (global clustering coefficient), unweighted mean shortest path length, and edge density. We tested whether observed network topology was consistent with features commonly seen in biological networks by comparing against 100 random graphs generated from a configuration model that preserved the observed degree distribution of the real network. The small-world coefficient sigma was computed as the ratio of the normalized clustering coefficient to the normalized path length (Humphries and Gurney, 2008): σ = (Clustering_obs_/Clustering_null_) / (Path length_obs_/Path length_nul_l). We used k-shell decomposition to characterize the core-periphery structure of the network (Carmi et al. 2007). The k-shell index of a gene is the largest value of k for which that gene is retained in the maximal subgraph in which every gene has at least k connections. Genes with degree less than k are removed, and the procedure is repeated until no genes remain. Genes removed in early rounds occupy the network periphery, whereas genes retained until later rounds occupy the network core. K-shell values were summarized by community and compared across communities.

### Community detection and resolution selection

Communities were identified using the Leiden algorithm (Traag et al. 2019) with the modularity objective function, implemented in igraph. To select an appropriate resolution parameter, we adapted the multi-resolution approach of Palma-Martínez et al. (2024), evaluating community structure across 50 resolution values in logarithmic space from 0.1 to 2.0. At each resolution, we retained the best partition from 10 independent Leiden runs, each with 20 iterations. These 20 iterations were used to assess the stability of communities for a given resolution by computing the mean pairwise Adjusted Rand Index (ARI).

We selected a resolution value of 1.0 because it was a value with both high modularity (0.289) and stability (ARI > 0.75 across the 0.6-1.2 range) where the largest number of differentiated major communities had emerged. At this resolution, 9 communities with 30 or more genes were identified for downstream analysis. The partition with the highest modularity across all runs at resolution 1.0 was used for all downstream characterization. To further validate that the observed communities reflect genuine network structure, the modularity of the resolution 1.0 partition (0.289) was compared against modularity obtained by running Leiden at the same resolution on 100 configuration model null networks preserving the observed degree sequence (0.100 ± 0.001).

### Community membership stability

The robustness of individual gene community assignments was assessed at the selected resolution (1.0) by running Leiden 50 times and aligning all partitions to the enrichment-run reference partition by maximum-overlap matching, following the approach of Palma-Martínez et al. (2024). For each gene, proportional membership was computed as the fraction of runs assigning that gene to each community. Genes were classified as “admixed” if no single community captured more than 80% of their assignments across the 50 runs. Per-community stability was summarized as the mean self-assignment proportion and the fraction of admixed genes. Results were visualized as a proportional membership plot with genes sorted by dominant community and stability, analogous to admixture plots in population genetic structure analysis. To visualize gene membership and community emergence and fragmentation across the resolution landscape, all partitions were aligned to a resolution 1.0 reference by maximum-overlap greedy matching of community labels in a resolution plot (Palma-Martínez et al. 2024).

### Cis-chromosomal edge composition

To distinguish gene functional-state covariance from shared deletions of physically linked genes, the distance composition of intra-community edges was assessed for each major community. Using the pairwise physical distance between genes from the EMMAX output, edges were classified as trans-chromosomal, cis-chromosomal but distant (>10 kb), or linked (<10 kb). The proportion of each edge class was computed per community.

### Functional enrichment

Functional enrichment of network communities were assessed with three complementary annotation sources accessed from TAIR: Gene Ontology terms (Biological Process and Molecular Function; Araport11, accessed April 22, 2025), InterPro protein domains (InterPro v. 99, March, 2024), and the gene functional descriptions supplied by TAIR for Araport11 (accessed June 30, 2025). GO enrichment was carried out using the topGO R package (Alexa and Rahnenfuhrer, 2023) with the weight01 algorithm. This algorithm accounts for hierarchical structure in the GO graph by conditioning a term’s test result on the results of neighboring terms to reduce redundancy among related terms and accounting for non-independence of GO terms. We follow the topGO authors recommendation of not performing multiple test correction on the p-values produced by the weight01 algorithm. For each network community, we performed two tests. First, a Fisher’s exact test for overrepresentation of a community against a background gene set. Second, a Kolmogorov-Smirnov test which uses the within-community degree of a gene as a gene-level score, which acts as a degree-weighted enrichment test that prioritizes functions whose annotated genes are in highly connected positions within a community. Terms significantly overrepresented by both the Fisher’s test (p < 0.05) and KS (p < 0.05) were designated as “core” functions of a community.

InterPro domain enrichment was assessed using a hypergeometric test for each community against a background set of genes with the Benjamini-Hochberg FDR correction. Centrality of the protein domains in a community was determined using a one-sided Wilcoxon rank-sum test comparing within-community degree of domain-carrying and non-carrying genes. Enrichment of functions from gene descriptions were performed as above, using a set of keyword searches against the Araport11 gene descriptions. All enrichment tests were performed using two background gene sets: all annotated Arabidopsis genes (global) and all genes within the LoF covariance network (network). For all annotation types, terms were classified by distinctiveness based on the number of communities in which they were significantly enriched.

### Network visualization

The global covariance network was visualized in Gephi (Bastian et al. 2009). Node and edge tables were exported from igraph with each gene annotated by its Leiden community assignment, k-shell value, and degree. For legibility, the visualized network was restricted to the strongest 25% of edges by association strength, and nodes were colored by community membership. Layout was computed using the ForceAtlas2 algorithm. This filtering was applied for visualization only; all quantitative analyses used the complete network.

### Adaptive loss-of-function enrichment

To test whether communities were differentially enriched for genes with prior evidence of adaptive loss of function, we cross-referenced network genes against the gene set analyzed by Monroe et al. (2018), who tested 2,088 *Arabidopsis* genes for associations between LoF genotype and flowering time, drought escape timing, latitude, and mean annual temperature. Genes with nominal association p < 0.05 for each variable were extracted from their Supplementary File 3. For each community and each variable, we tested for enrichment with a hypergeometric test comparing the number of genes in a community that showed a significant association to the expected number given the community size. We repeated this analysis using only genes whose associations remained significant after Bonferroni correction in the original study.

### Inter-community analysis

To assess network connectivity between modules, we computed an inter-community edge density for all pairs of major communities (≥30 genes) by dividing the total number of observed edges between a community pair by the number of possible edges between that pair and compared it against a null expectation generated by permuting community labels 1,000 times while preserving the network topology. To compare inter-community connectivity across community pairs of different sizes, a mixing parameter was computed as the inter-community edge density divided by the geometric mean of the two communities’ intra-community densities.The mixing parameter distance matrix was visualized as a heatmap with hierarchical clustering.

### Community LoF burden and population structure

Community-level gene content variation was quantified at the accession level by calculating community LoF burden for each accession as the proportion of genes assigned to a given community carrying a loss-of-function state. Burdens were calculated for the four largest communities across all 1,012 Eurasian accessions. Pairwise Pearson correlations among community burdens were computed to assess whether communities covaried across accessions. To rule out population structure, we controlled for the recorded latitude and longitude of an accession using the ppcor R package (Kim 2015).

Associations between community burden and geography were assessed by Pearson correlation with latitude and longitude. Differences among admixture groups (Alonso-Blanco et al. 2016) were assessed by one-way ANOVA, with the proportion of variance explained reported as R². Accession scores on the first five principal components from the Eurasian accession PCA were correlated with community LoF burden to assess whether communities occupied distinct positions along the major axes of gene content variation.

## Results

### Functional gene content variation in the Arabidopsis dispensable genome

To characterize gene content variation across *Arabidopsis thaliana*, we used a recently published catalog of 91,751 naturally occurring loss-of-function variants identified from whole-genome sequencing data of 1,135 accessions in the 1001 Genomes collection (Zhao et al. 2026). This catalog integrates SNP-based LoF variants (frameshift mutations, premature stop codons, splice site disruptions) with structural variant deletions, all called relative to the Columbia (Col-0) reference genome. Independent LoF alleles affecting the same gene were collapsed into a binary functional state, producing a matrix of 12,182 genes with segregating loss-of-function across 1,112 accessions. These inferred gene losses are supported by reduced expression of LoF alleles relative to functional counterparts and are strongly predictive of gene dispensability in a 32-accession Arabidopsis pangenome (Zhao et al. 2026), supporting their use as a proxy for gene function presence-absence variation. As this approach identifies gene function loss relative to the Col-0 reference, it will undercount LoF alleles in genes absent from or already nonfunctional in Col-0.

Gene loss varied substantially across accessions (median 1,997 affected genes per accession, range 676-2,751) and differed among admixture groups (Figure 1A-C). Principal component analysis of gene content variation revealed that the first axis (PC1, 5.2% of variance) separated North American from Eurasian accessions, reflecting a known colonization bottleneck of the HPG1 lineage (Figure 1D; Exposito-Alonso et al. 2018). We find that this bottleneck fixed over 1,500 gene losses in the founding population, leading to extremely low variation in gene-loss profiles for the entire HPG1 population. Despite this, the HPG1 gene content profile was no more deleterious than typical Eurasian accessions across measures of selective constraint, expression breadth, and lethality (Figure S1), suggesting that even demographically driven gene loss operates within functional constraints. The HPG1 bottleneck demonstrates that demographic processes can generate extensive correlated gene content variation, and we exclude this population from downstream analyses to alleviate this strong demographic signal.

### Genes pairs with associated LoFs are enriched for shared function

Having established that demographic processes can generate extensive gene functional-state covariance, we next asked whether non-random associations among dispensable genes persist after controlling for population structure and shared ancestry. We excluded North American accessions and carried out an all-by-all pairwise association testing using the EMMAX mixed linear model (Kang et al. 2010) on the remaining 1,012 Eurasian accessions across 12,182 genes, treating each gene’s binary LoF status as a phenotype and testing all other genes as predictors while controlling for kinship. After filtering for FDR-adjusted p < 0.05, requiring both genes in each pair to have LoF frequency between 0.05 and 0.95, and averaging reciprocal tests to yield unique gene pairs, we identified 150,033 significant associations among 5,679 genes from an initial 148 million pairwise tests. We refer to these relationships as LoF associations: positive associations indicate that loss-of-function states tend to occur in the same accessions, whereas negative associations indicate that the loss-of-function state at one gene is less frequent among accessions carrying loss-of-function at the other. The majority of LoF associations were positive rather than negative (140,568 positive associations, among 5,673 genes vs 9,465 negative associations among 4,528 genes), and most genes (4,522) participated in both positive and negative associations.

Both positively and negatively associated gene pairs were enriched for closely linked genes relative to the genomic null expectation. Compared to a null expectation of 0.02% of all possible gene pairs falling within 10kb (the approximate distance at which LD decays by 50% in Arabidopsis; Kim et al. 2007), 7% of positively associated gene pairs and 15% of negatively associated gene pairs were within 10kb. For positively associated gene pairs, this enrichment is consistent with tandem co-deletion through shared structural variants; in negatively associated pairs, it may reflect selection maintaining at least one functional copy of tandem duplicates. However, the majority of co-occurring pairs (74%) were trans-chromosomal, close to the genomic expectation of 79%. This trans-chromosomal covariance in gene functional state, maintained after controlling for kinship, argues against physical linkage and residual population structure as primary explanations for the observed covariance and is consistent with selection acting on multilocus combinations.

Following Whelan et al. 2021, we hypothesized that if LoF associations preferentially occur among functionally related genes, the associated loci should share biological function and expression patterns more than random pairs of genes. To test this, we compared pairwise coexpression and GO annotation overlap of significantly associated gene pairs against null expectations. For the enrichment analyses, we restricted to gene pairs where both genes were represented in the Klepikova developmental transcriptome atlas (Klepikova et al. 2016), retaining 117,386 co-occurring and 8,337 avoidant pairs. We created 100 permuted datasets in which gene pair assignments were independently shuffled, preserving the pool of genes while breaking the observed pairings. Coexpression was measured as Spearman correlation of expression profiles across all tissues in the Klepikova developmental transcriptome atlas (Klepikova et al. 2016). Functional annotation overlap was measured as the proportion of gene pairs sharing at least one GO Biological Process term as in Whelan et al. (2021). Co-occurring gene pairs showed significantly higher mean coexpression than all 100 permuted datasets with an observed mean coexpression of 0.048 vs a mean coexpression of 0.042 across 100 permutations (p< 0.01; Figure 2A). Co-occurring pairs also showed significantly greater overlap in GO Biological Process annotations than expected by chance (observed overlap 23.9% vs permuted mean 22.6%; p < 0.01; Figure 2B). Similar enrichment was found for GO Molecular Function and Cellular Component annotations (Figure S2). Avoidant gene pairs showed qualitatively similar enrichment for all metrics, (Figure S3). Together, these results indicate that genes with covarying functional states are enriched for similar biological function and expression, consistent with our hypothesis.

**Figure 2.**
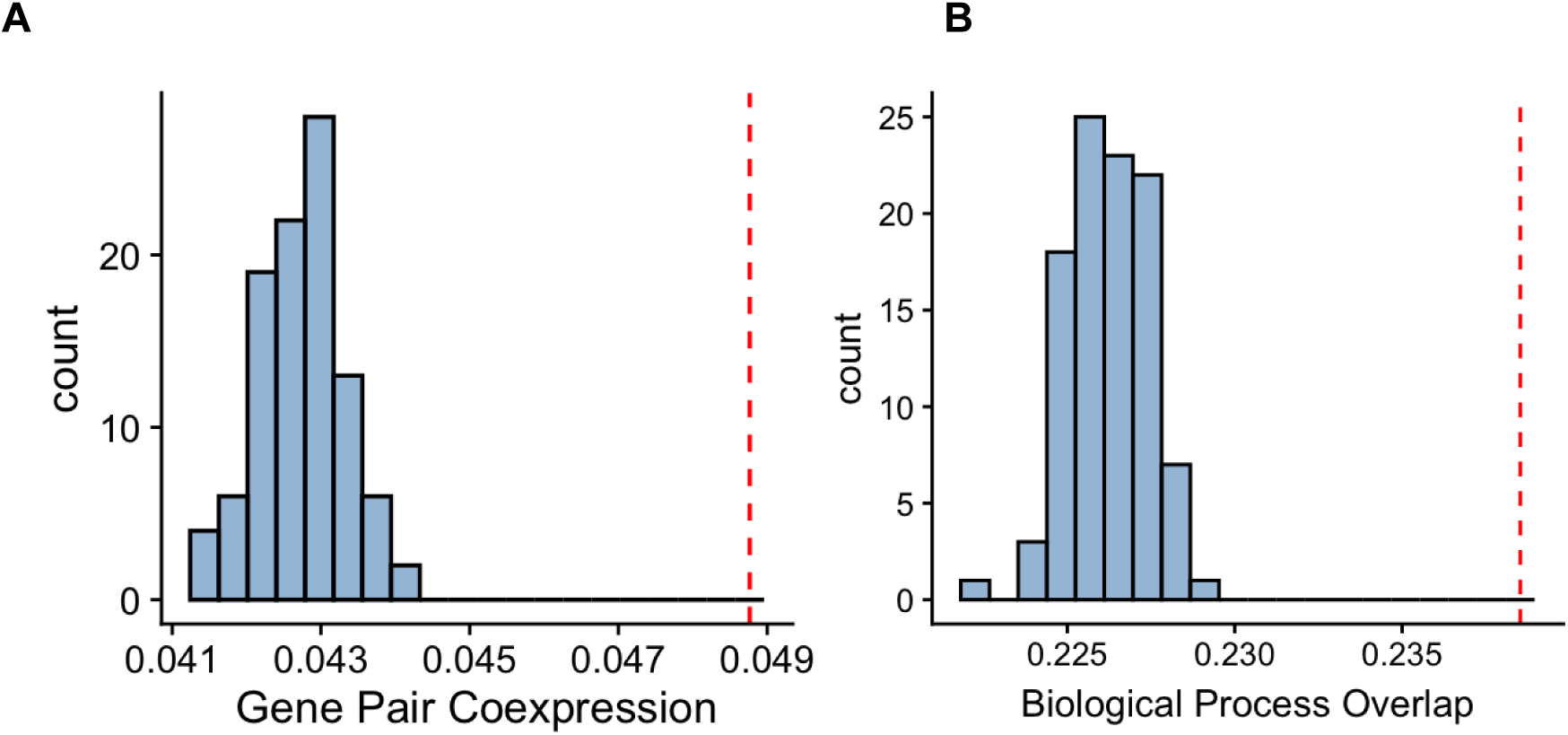
Functional similarity amongst genes with covarying functional states. A) Mean coexpression of gene pairs with significant positive covariation of functional states (red dashed line) compared to distribution of mean coexpression (Spearman’s r) of 100 permuted gene pairs (blue distribution). B) Mean overlap of biological process GO terms for genes pairs with significant positive covariation of functional states (red dashed line) compared to distribution of mean overlap of biological process GO terms of 100 permuted gene pairs (blue distribution)

### The functional-state covariance network has non-random organization

While the pairwise analysis established that genes with covarying LoF states are enriched for shared function, it is unable to reveal whether these associations are organized into a higher-order structure. To test this, we constructed a network from all significant positive LoF associations, representing each gene as a node and each association as a weighted edge based on EMMAX effect sizes. The resulting LoF covariance network contained 5,673 nodes and 140,568 edges. To better understand the topology of the network, we compared the LoF covariance network to 100 random networks with matched degree distributions in terms of their clustering coefficients (transitivity), a measure of the tendency for neighboring genes in a network to form interconnected groups, and the average shortest path between genes. We found that the observed network had substantially greater clustering than expected by chance (transitivity = 0.262 vs. 0.114 in degree-matched networks; p < 0.01), and maintained short paths between genes (observed mean shortest path = 2.99 vs. 2.54), suggesting that genes form tightly interconnected neighborhoods that remain connected to the larger network by a relatively small number of steps in the network, a hallmark of many biological networks (Humphries and Gurney, 2008; Table 1).

**Table 1.** Summary statistics of global network topology.

| Statistic | Observed | Expected |
| --- | --- | --- |
| Nodes (genes) | 5,673 | — |
| Edges (associations) | 140,568 | — |
| Edge density | 0.0087 | — |
| Mean degree | 49.6 | — |
| Clustering coefficient (transitivity) | 0.262 | 0.114 |
| Mean shortest path length | 2.99 | 2.54 |
| Small-world coefficient $\sigma$ | 1.95 | — |
| Modularity (Leiden, resolution 1.0) | 0.289 | 0.100 ± 0.001 |
| Communities with $\geq 30$ genes | 9 | — |

To better characterize the structure and content of these neighborhoods, we next queried whether the neighborhoods indicated by the clustering coefficient could be formally partitioned into discrete groups of genes with unusually dense internal connections. We used the Leiden algorithm to formally partition the network into “communities” of genes with dense internal connections and sparse external ones (Traag et al. 2019; Palma-Martínez et al. 2024; see methods). This community detection identified nine communities with 30 or more genes. Community sizes varied widely: C1 contained 3,134 genes (53% of the network), whereas the remaining communities ranged from 31 to 995 genes. (Fig. 3A-C).

**Figure 3.**
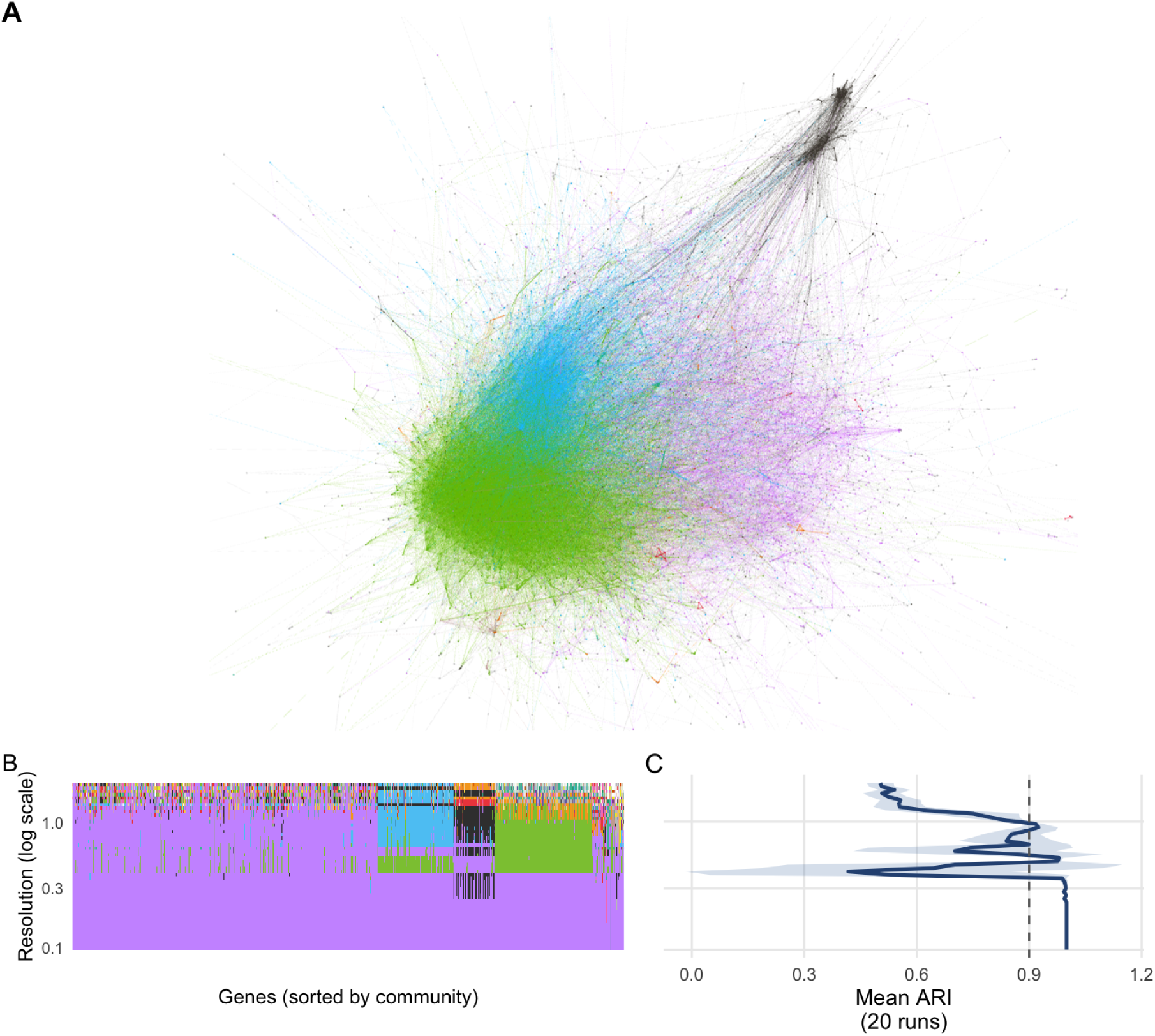
Higher-order structure of loss-of-function covariation. A) Global network of LoF covariation, filtered to only show the top 25% strongest edges for visualization purposes, visualized in Gephi. Colors represent community assignments from the optimal resolution value of 1: Purple is C1, Blue is C2, Black is C3, Green is C4, Orange is C5, Red is C6. Nodes are colored by their community membership. B). Resolution plot of gene community membership across different resolution parameters used in community detection. Each point along the x-axis is a single gene, with colors representing that gene’s community membership. The y-axis is the log-scaled resolution parameter for the leiden community detection algorithm. Higher parameter values generate smaller communities while lower values tend to generate larger communities. The plot visualizes the changes in a gene’s group assignment as the resolution value increases, showing the emergence and breakdown of distinct communities across parameter space. C). Adjusted Rand Index values quantifying mean community stability across 20 independent Leiden algorithm runs. ARI values closer to 1 indicate a higher proportion of genes falling into the same communities across independent runs. The mean stability of the 20 runs was plotted across the entire range of resolution values, showing changes in stability for different inferred communities.

### Network communities are topologically and functionally distinct

Although nine communities were identified, they differed markedly in their stability and position within the network. To assess community stability, we assessed the proportion of runs assigning each gene to the same community across 50 independent Leiden runs to measure the reproducibility of each community. Four large communities (C1 with 3,134 genes, C2 with 784 genes, C3 with 428 genes, and C4 with 995 genes) and two small communities (C10 with 39 genes and C11 with 31 genes) were highly stable, with genes assigned to the same community in over 75% of runs on average. The other communities (C5 with 90 genes, C6 with 61 genes, and C8 with 43 genes) showed substantially lower assignment stability (<65%; Figure 4A). We next used k-shell decomposition (Carmi et al. 2007) to explore the layers of network connectivity by iteratively removing genes with few connections. Genes with higher k-shell scores remain until later rounds and occupy the network core, whereas genes removed early are more peripheral. Two of the stable communities (C2 and C4) were substantially more deeply embedded in the network than the other communities (median k-shell values 37 and 64 versus 10-15 for the other communities; Figure 4B).

**Figure 4.**
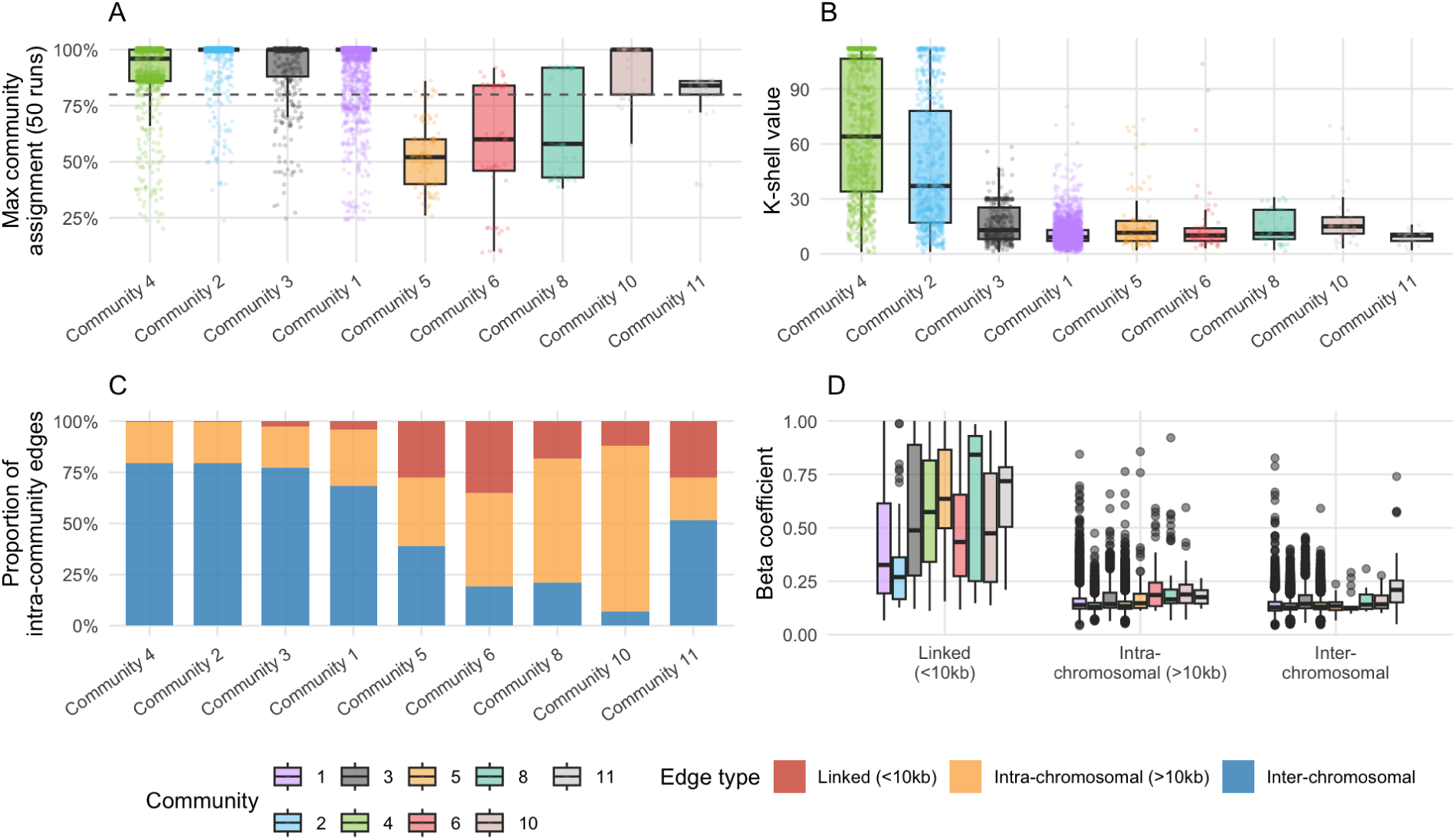
Topological characterization of network communities. A) Distribution of the proportion of 50 replicated runs in which a gene is assigned to the same community for all communities with > 30 genes. B) Distribution of gene k-shell value in the network, for each community with > 30 genes. C). Breakdown of the proportion of edges deriving from gene pairs in physical linkage (Red), unlinked genes on the same chromosome (yellow), and gene pairs located on different chromosomes (blue). D). Distribution of the strength of association between gene pairs for pairs that are physically linked, unlinked but on the same chromosome, and on different chromosomes.

The genomic composition of community edges further distinguished these network structures. The four large stable communities (C1-C4), were dominated by inter-chromosomal associations (68-80%) while only 0.4-4.2% of edges connected genes expected to be in physical linkage (<10 kb apart; Figure 4C). Thus, C3 resembled the deeply embedded C2 and C4 modules in being primarily comprised of distributed, genome-wide associations, despite its more isolated network position. C1 was also predominantly trans-chromosomal (68%), but contained a larger contribution from distant intra-chromosomal associations (27.4%) and linked pairs (4.2%). In contrast, the smaller communities contained substantially more physically linked associations (12–35% of internal edges) and were more dominated by both linked and distal intra-chromosomal connections, which together accounted for 39–93% of internal edges overall. Physically linked pairs also had substantially larger association coefficients than distant intra– or inter-chromosomal pairs (median β = 0.42–0.58 vs. 0.13–0.16; Figure 4D). This indicates that the smaller communities are more driven by local structural variation while the larger communities were defined primarily by distributed associations among unlinked genes.

Network communities also showed distinct functional compositions. We tested network communities for functional enrichment and compared enrichments across communities. Each community was functionally characterized using GO terms, InterPro protein domains, and TAIR functional descriptions, with enrichment assessed against both genome-wide and network-specific backgrounds. The within-community degree of a gene was additionally used to prioritize functions concentrated among highly connected genes in a community. Functions supported by both overrepresentation and degree-weighted enrichment were designated as core functions.

Across these sources of evidence, the four large stable communities showed distinct functional signatures that converged. C4 (995 genes) was strongly enriched for NB-ARC and TIR domains as well as defense response and signal transduction, consistent with NLR-mediated intracellular immune perception. C3 (428 genes) was enriched for structural constituents of the cell wall, glutathione metabolism, cation exchangers, and wall-associated receptor kinases, consistent with a role in cell-wall and apoplastic homeostasis. C2 (784 genes) showed enrichment for ADP binding and leucine-rich repeat proteins lacking an NB-ARC domain, and signal transduction and stress response were concentrated among highly connected genes in the community. Distinctive enrichments in serpins and other protease inhibitors, together with cullin scaffolds, suggest it is a stress-responsive regulatory and signaling module. C1, the largest community (3,134 genes), showed enrichment for multiple specialized metabolism processes, including diterpenoid and sesquiterpenoid biosynthesis, together with cytochrome P450, Gnk2-like domains, and nitrilases. These functions were concentrated among more centrally connected genes, whereas F-box and ubiquitin-associated genes were less connected in the community.

Among the smaller, more peripheral communities, C5 was characterized by enrichment for SAUR (small auxin-up RNA) proteins, immune-associated GTPases and secretory/reproductive functions; C6 by NAC and S-domain receptor kinase signatures, including S-locus-associated domains, consistent with pollen or self/non-self recognition and signaling; C8 by synergid death, double fertilization, and miRNA catabolic process and binding; C10 by RNA-directed DNA methylation; and C11 by cell wall loosening and expansin-associated functions (Table 2; Dataset S1). While enrichment of defense and stress-related genes in the dispensable genome is well documented in plant pangenomes, these results indicate that the dispensable genome partitions defense-associated functions into distinct network communities corresponding to different layers of biological organization.

**Table 2.** Summary of network community functional enrichment.

| Community | N genes | Functional identity | Stability (%) | Key enrichments |
| --- | --- | --- | --- | --- |
| C1 | 3,134 | Specialized defense metabolism | 95.6 | GO-BP, InterPro, Description (core: GnK2, terpene synthases, CYP450, nitrilase) |
| C2 | 784 | Stress signaling / proteolytic regulation | 95.9 | GO-BP, GO-MF, InterPro, Description (core: signal transduction, serpin, cullin, protease inhibitor, stress response) |
| C3 | 428 | Cell wall / apoplastic homeostasis | 89.5 | GO-MF, InterPro, Description (core: cation exchanger, germin, glutathione, RLK) |
| C4 | 995 | NLR / intracellular perception | 89.7 | GO-MF, GO-BP, InterPro, Description (core: NB-ARC, APAF, TIR NADase, LRR) |
| C5 | 90 | Immune GTPase / secretory signaling | 47.6 | InterPro (core: SAUR, AIG1 GTPase) |
| C6 | 61 | Self/non-self recognition | 56.9 | InterPro, GO-MF (core: NAC domain, S-locus glycoprotein, calmodulin binding) |
| C8 | 43 | Gametophyte / RNA regulation | 64.3 | GO-BP (enriched: synergid death, double fertilization, miRNA) |
| C10 | 39 | RNA-directed DNA methylation | 91.9 | GO-BP (enriched: RNA-directed DNA methylation) |
| C11 | 31 | Cell expansion / stress physiology | 77.8 | GO-MF, InterPro (core: sulfotransferase; enriched: expansin, cell wall loosening) |

### Inter-community connectivity reveals evolutionary coupling between functional modules

The identification of distinct modules allows us to investigate the higher-order structure of the LoF covariance network by characterizing the connectedness between different network communities. We computed a mixing parameter for each community pair, defined as the inter-community edge density normalized by the geometric mean of the two communities’ intra-community densities. Hierarchical clustering of the resulting matrix revealed structured connectivity between network communities (Figure 5). The strongest association in the network was between C2 and C4 (mixing = 0.403), with additional strong connections between C1 and C4 (0.182) and C1 and C2 (0.175), identifying C1–C2–C4 as the principal interconnected component of the network. This core component, anchored by the NLR-enriched C4 community, showed broad connectivity to C5, C6, C8, and C10. These communities showed weak connectivity to one another but preferential connections to the larger network, consistent with their functioning as peripheral interface communities rather than autonomous modules. C3, on the other hand, showed moderate connections to C1 and C2 (0.111 and 0.092) but weak connectivity to C4 and the smaller communities, consistent with being an autonomous functional module. C11 was nearly isolated from the remainder of the network, suggesting that it is a functional component largely independent of the defense axis.

**Figure 5.**
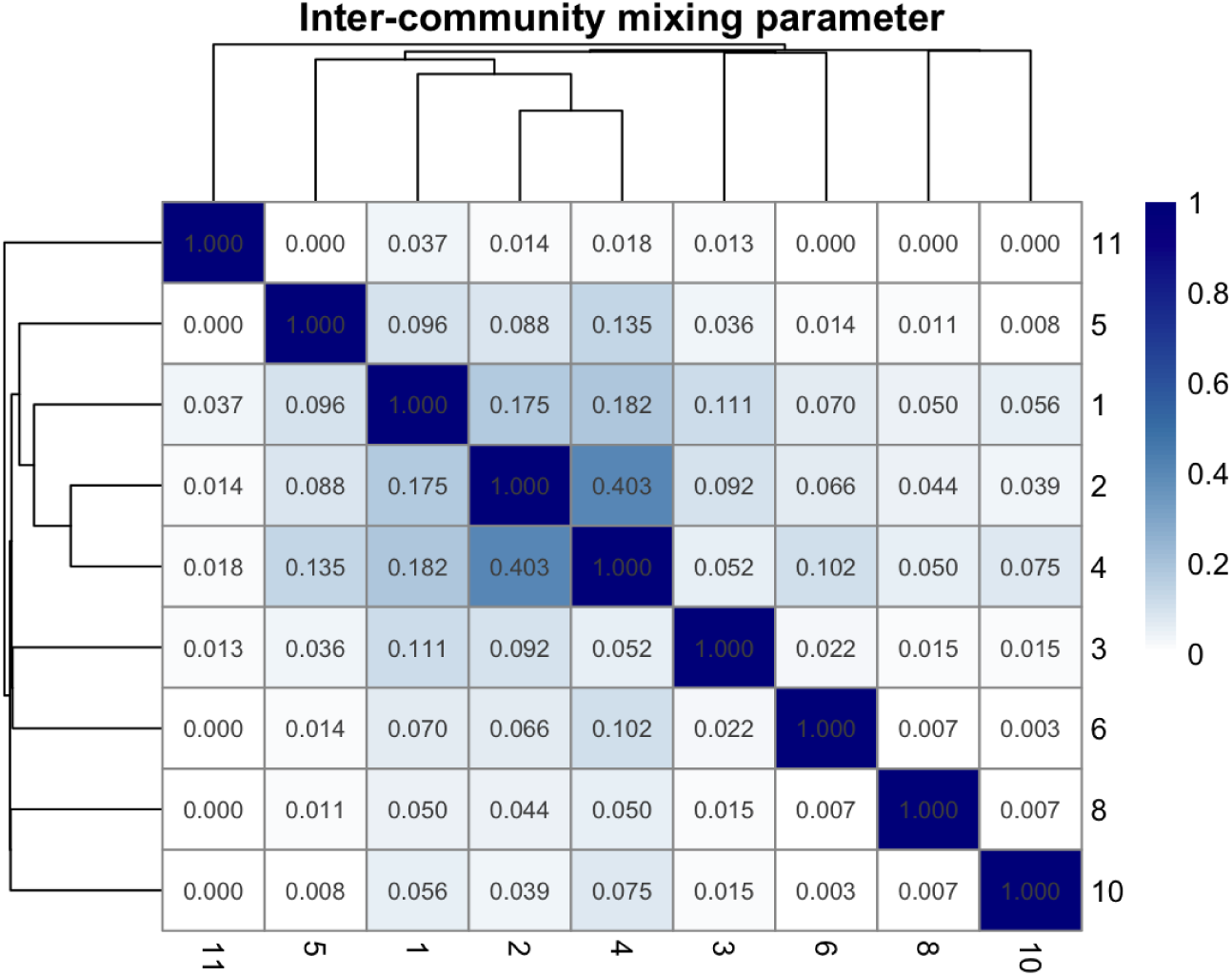
Intercommunity structure in the global covariance network. Mixing parameters for each pair of communities, defined as the inter-community edge density normalized by the geometric mean of the two communities’ intra-community densities. Rows and columns are ordered by hierarchical clustering (complete linkage) on the mixing parameter matrix. Higher values indicate that two communities are more connected to one another than expected given their internal densities.

**Figure 6.**
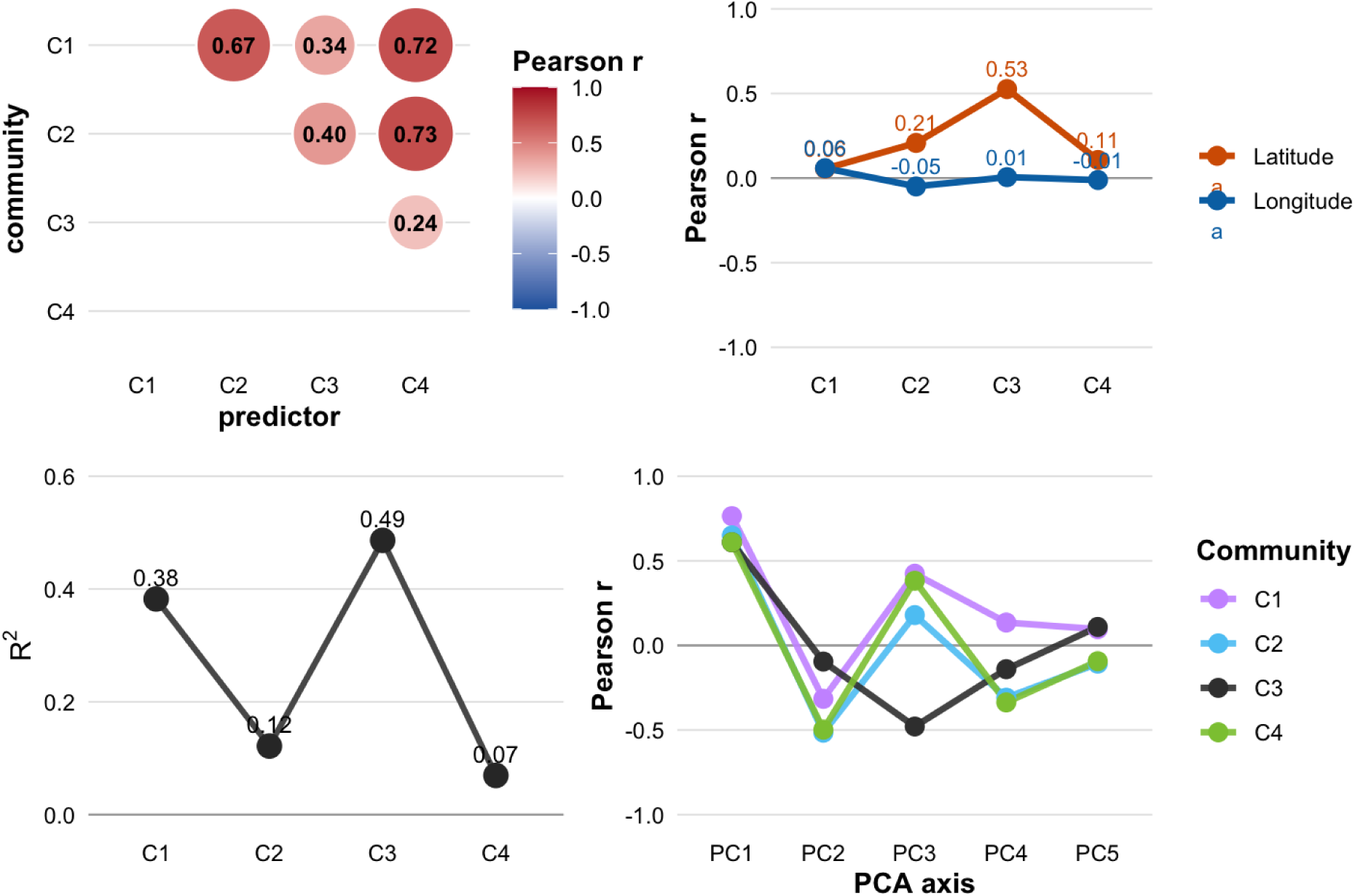
Community LoF burden across accessions and relationship to population structure. For each accession, community LoF burden was calculated as the proportion of genes assigned to a community carrying a loss-of-function state. (A) Pairwise Pearson correlations of community LoF burden across 1,012 Eurasian *Arabidopsis thaliana* accessions. (B) Correlations between community LoF burden and geographic coordinates. (C) Proportion of variance in community LoF burden explained by admixture group membership (one-way ANOVA R^2^). (D) Correlations between community LoF burden and accession scores on the first five principal components of the Eurasian LoF matrix.

This pattern of connectivity between modules suggests that the covariance network may reflect not only independent modules but also the functional interactions between distinct biological processes. We next asked whether the communities occupying different positions in the network also differed in their evolutionary history. Monroe et al. (2018) identified 2,088 Arabidopsis genes carrying putatively adaptive loss-of-function alleles associated with flowering time, drought timing, latitude, and temperature. For each environmental or phenotypic variable, we tested for community enrichment of genes with nominal association (p < 0.05 in the published analysis) against the distribution of those genes within the network. Cross-referencing these adaptively evolving genes with our network revealed that C2 was significantly enriched (33% observed vs 26% expected, p = 3.2 × 10^−6^) and specifically those associated with flowering time (p = 7.8 × 10^−7^), drought timing (p = 1.0 × 10^−4^), and latitude (p = 2.6 × 10^−8^). In contrast, C4 showed no significant enrichment for these loci. These enrichments remained significant when restricted to the more stringent LoF-climate associations that are significant after Bonferroni correction. Together, these results identify C2 as a network community enriched for genes associated with environmental adaptation, particularly flowering time, drought response, and geographic variation, consistent with the earlier finding that C2 most likely functions as a regulatory and signaling module tied to stress responses. Because C2 is also the community most strongly connected to the NLR module C4, these results suggest that environmental stress response and pathogen defense are coupled within the functional-state covariance architecture.

The evolutionary distinction between communities was also evident at the level of accession-wide gene content. For each accession, a community LoF burden was calculated as the proportion of genes assigned to a community that carried a LoF state. Community LoF burdens were correlated across accessions, with the strongest correspondence between C2 and C4 (r = 0.728), mirroring their strong inter-community network affinity (Figure 6A). These communities showed weak associations with latitude (r = 0.21 and 0.11, respectively) and longitude (r = –0.05 and –0.01, respectively), and their intercommunity correlation was nearly unchanged after controlling for latitude and longitude (partial r = 0.727; Table S1; Figure 6B). In contrast, C3 was less strongly correlated with the other major communities and showed the strongest geographic differentiation, with community LoF burden increasing with latitude (r = 0.526). Community burden also differed among admixture groups (Figure 6C), particularly for C3 (R^2^ = 0.486) and C1 (R^2^ = 0.382). PCA of the Eurasian LoF matrix further showed that the stable communities occupy distinct positions along several major axes of gene-content variation (Figure 6D). C2 and C4 had similar correlations with the first PCs, whereas C3 differed most strongly along PC3 and PC5, consistent with its distinct geographic and population-genetic distribution. Together, these analyses show that the modular structure of the covariance network is reflected not just in network topology and function, but in how evolutionary processes shape functional gene content and its distribution across natural populations.

## Discussion

Pangenomic analyses have firmly established that genomes of many species vary substantially in their gene content. These studies have conventionally treated dispensable genes as independent loci with unrelated evolutionary histories. Instead, our results suggest that this view of pangenomic variation is insufficient. Across our 1,012 Eurasian Arabidopsis accessions, the functional states of genes showed extensive covariance, after accounting for kinship and population structure. These associations were predominantly between unlinked loci on different chromosomes, were enriched among gene pairs with shared expression profiles and biological functions, and resolved into a higher-order covariance network comprising distinct functional communities. Thus, gene content variation in Arabidopsis is not simply a ledger of independently segregating dispensable genes, but is organized into interacting sets of functional modules.

These results build on previous observations from microbial pangenomics, in which *Pseudomonas* (Whelan et al. 2021) and *E. coli* (Hall et al. 2021; Beavan et al. 2024) accessory genes showed non-random patterns of co-occurrence driven by functional relationships among genes. The observation of higher-order organization in a eukaryotic pangenome, despite different genetic mechanisms enabling gene content variation than in prokaryotes, suggests that the functional interaction of genes is a general factor driving pangenome structure. Our results also match a core prediction of multilocus population genetic theory: that when selection acts on combinations of loci, non-random associations are expected to emerge, including among physically unlinked loci. The functionally coherent network communities driven by unlinked loci are consistent with multilocus assemblages and therefore provide evidence that selection is acting on multilocus combinations rather than independently on genes.

While we provide strong evidence for the existence of these assemblages, we cannot yet distinguish the mechanism(s) that generate them. The multilocus covariance we find can arise from epistatic interactions among functionally complementary loci, as in coadapted gene complexes (Lewontin and Kojima, 1960), or from polygenic adaptation in which multiple loci respond to shared selective pressures (Latta, 1998). Distinguishing the role of these two processes will require experimental or ecological data that can resolve the fitness consequences or evolutionary trajectories of specific multilocus combinations. It is notable, however, that the biological processes most prominently represented in our network are among those for which epistasis is best documented in Arabidopsis. Combinations of NLR alleles carried by different accessions can produce autoimmune phenotypes with severe fitness costs, such that the fitness effect of an NLR depends on which other NLRs are present in the same genome (Chae et al. 2014; Lee and Chae 2025). Natural variation in specialized metabolism is likewise shaped by epistatic interactions among loci (Chan et al. 2010; Rowe et al. 2008) and genetic incompatibilities acting through pollen accumulate rapidly between diverging *Arabidopsis* populations (Condon et al. 2025). That the communities we recover are enriched for precisely these functions is highly consistent with epistatic selection contributing to their coherence, though it does not establish it.

### A systems biology perspective of pangenomic variation

Our analysis also illustrates a reciprocal relationship between systems biology and evolutionary genetics. Systems biology has long contended that natural variation is best interpretable in the context of the networks and pathways in which genes operate (Kliebenstein, 2010; Ogura and Busch, 2016). What distinguishes the approach taken here is its scale and independence from prior pathway specification. The work dissecting flowering time and glucosinolate biosynthesis used natural variation to probe characterized mechanisms within known pathways already identified as candidates. With this approach, the covariance structure of gene functional state across the genome recovers functional organization without pre-specifying pathways or measuring phenotypes directly. The network approach produces communities that are empirical representations of gene functions that tend to be retained or lost together in natural genotypes. The resulting communities can then be interpreted against known biology or used to generate novel and testable hypotheses. The distinct topologies of communities and their patterns of interconnection provide further information about how these functional systems are organized within the genome and how their components may respond collectively to ecological pressures. This is most apparent in the relationship between C4, dominated by rapidly evolving NLR genes, and C2, enriched for previously identified adaptive LoF loci associated with flowering time, drought timing, and latitude.

In our covariation network, the C2 and C4 communities represented the two most deeply embedded and interconnected communities. The functional enrichments we recovered for these communities, and their connection in the covariance network, strongly suggests that environmental and developmental adaptation and intracellular pathogen perception are coupled at the level of pangenome functional-state variation. Such a coupling is ecologically plausible. Across Europe, phyllosphere composition covaried with latitude and drought conditions, and drought conditions in conjunction with natural variation in *Arabidopsis* shape the bacterial communities colonizing leaves, a clear example of abiotic environments shaping the biotic interactions a plant experiences (Karasov et al. 2024). At the molecular level, immune signaling also intersects extensively with developmental and environmental-response pathways, including regulatory networks influenced by NLR signaling (Ariga et al. 2017; Hu et al. 2026; Wang et al. 2011; Zarattini et al. 2021). Our results demonstrate that this functional integration at the cellular and molecular level shapes the composition of the dispensable genome. Critically, the fact that the interconnectivity between C2 and C4 is strongest topological feature in the global network appears consistent with recent theorizing that growth, development, environmental response, and immunity are not necessarily opposing evolutionary axes, but instead are better understood as an integrated system, coordinated as plants evolve in response to dynamic ecological pressures (Kliebenstein, 2016).

C1 provides an even clearer example of how network architecture can reveal biological relationships that are otherwise cryptic. This community contained multiple specialized-metabolism pathways and acted as a third component in the integrated network core along with the NLR immune perception and environmental response. This topology is consistent with an integrated role for specialized metabolism and the accumulating evidence that specialized metabolites are multifunctional compounds simultaneously involving defensive compounds, regulators, and components of primary metabolism (Erb and Kliebenstein, 2020). Indeed, the most central genes in this community were enriched for well-established multi-functional pathways. This includes metabolism and regulation of terpenoids, a diverse family of compounds involved in defense, herbivore and pathogen interactions, pollinator attraction and other ecological interactions (Huang et al. 2010; Kang et al. 2025; Tholl and Lee, 2011); Nitrilases and nitrile-specifier proteins, linking metabolism of indole-derived glucosinolate defense compounds to auxin and downstream development and stress response (Bartling et al. 1994; Cackett et al. 2022; Hornbacher et al. 2022; Normanly et al. 1997; Wójcikowska, 2026); and brassinosteroid biosynthesis and homeostasis, a plant hormone known to regulate growth, development, and stress responses (Clouse and Sasse, 1998; Nolan et al., 2020). The position of C1 between the environmentally associated C2 and immune-associated C4 communities is striking in this context, suggesting that the C1–C2–C4 architecture reflects the integration of environmental sensing, metabolism, and immunity within a shared evolutionary landscape.

The remaining communities revealed additional architectural modes that suggest a rich diversity of topologies among network communities. The smaller communities had weak intra-community connectivity, often containing disjoint sets of genes that were only connected indirectly via other communities. They also showed weak connectivity to other small communities. Instead, their defining feature was the preferential connections to the larger network, and enrichment for functions associated with recognition, receptor signaling, regulation, secretion, and reproduction. The position of these communities indicates that some covariance network communities may be defined more by their larger core functional systems, like immunity and environmental response, rather than as their own independent co-evolutionary modules. The enrichment of receptor and recognition functions in these peripheral communities is consistent with this integrative role, although the size and contribution of physically linked associations prevent more robust interpretation. C3 provides a final contrast in that it formed a stable, predominantly trans-chromosomal community that was relatively autonomous from the defense-centered network, with cell-wall and apoplastic functions providing a coherent biological interpretation of its network position and indicating that not all aspects of the dispensable genome are mediated through interactions with biotic and abiotic response systems.

### Limitations and Future Directions

Although our approach is powerful and provides strong evidence for higher-order structure of a complex eukaryotic pangenome, future work is needed to assess the generality of these results across eukaryotes. It is unclear how the selfing nature of *Arabidopsis thaliana* might affect our association approach or the interpretation of associations, and additional population-scale pangenomic studies are needed to replicate our results beyond this species. Additionally, the use of pangenome graph-based alignment would address the reference bias in this study, which prevented us from identifying well-known examples like FLC or AOP and MAM associations, as those are pseudogenes in the Col-0 reference genome. To better understand the role of functional interactions and the relative contribution of epistatic vs polygenic selection, functional validation with large-scale CRISPR arrays, as well as integration with other systems biology network approaches, like gene regulatory networks, protein-protein interaction networks and metabolic networks will be needed. There are likewise exciting parallels being developed, from single-cell RNAseq coexpression to gene expression programs, that may be complementary to the one explored here (Parker and Pennell, 2025).

## Conclusion

Our work shows that treating the natural occurrence of thousands of gene knockouts across diverse genotypes as functional perturbations and examining their covariance across diverse natural genotypes can reveal the higher-order organization of the dispensable genome. The correspondence between covariance structure, network topology, and biological function suggests that this approach provides a population-level readout of functional dependencies among genes and the interactions of the biological systems in which they operate. When analyzed in this way, patterns of natural variation in gene content can be a valuable resource for discovering the structure of biological systems.

## Competing Interests

The author(s) declare no conflicts of interest.

## Author Contributions

KAB, DJK, and JGM conceptualized the project; KAB and KZ performed analyses; KAB wrote the primary draft; KAB, DJK, KZ, and JGM edited the manuscript.

## Data availability

All data analysed in this study are publicly available. Loss-of-function variants were called from the 1001 Genomes Project SNP and indel release annotated with SnpEff (https://1001genomes.org/data/GMI-MPI/releases/v3.1/1001genomes_snpeff_v3.1/) and from the structural variant catalogue of Göktay et al. (2021). The kinship matrix was derived from the 1001 Genomes imputed SNP matrix (https://1001genomes.org/data/GMI-MPI/releases/v3.1/SNP_matrix_imputed_hdf5/). The collapsed loss-of-function allele matrix is described in and available from Zhao et al. (2026). Expression data are from the Klepikova et al. (2016) developmental transcriptome atlas. Gene Ontology annotations, InterPro domain assignments and gene functional descriptions were obtained from TAIR for Araport11. Previously identified adaptive loss-of-function associations were taken from Supplementary File 3 of Monroe et al. (2018). Complete functional enrichment results for all communities are provided as Dataset S1.

## Code availability

Code for the core analyses can be found at https://github.com/KevinABird/LoFNetwork

## Acknowledgements

This work was supported by the National Science Foundation Division of Integrative Organismal Systems grant 2208944 to KAB. We would also like to thank Dr. Nicholas Provart and the Bio-Analytic Resource for Plant Biology (BAR) for providing access to the Klepikova Arabidopsis expression atlas data.

## Notes

### Competing Interest Statement

The authors have declared no competing interest.

