## Supplementary material for "Gene functional-state covariance reveals modular structure in the *Arabidopsis thaliana* dispensable genome": Figure S1

**Figure S1. Gene content of the North American HPG1 lineage compared with Eurasian accessions.** Distributions show, across all Eurasian accessions, the mean value of each measure taken over the set of genes carrying a loss-of-function state in that accession: (A) the ratio of non-synonymous to synonymous substitution rates (dN/dS), (B) the neutrality index, (C) tissue breadth of expression, and (D) the proportion of genes whose knockout is reported as lethal. Red dashed lines show the corresponding values for the set of genes fixed for loss of function within HPG1. Fixed HPG1 losses fall within the range of standing variation for all four measures.

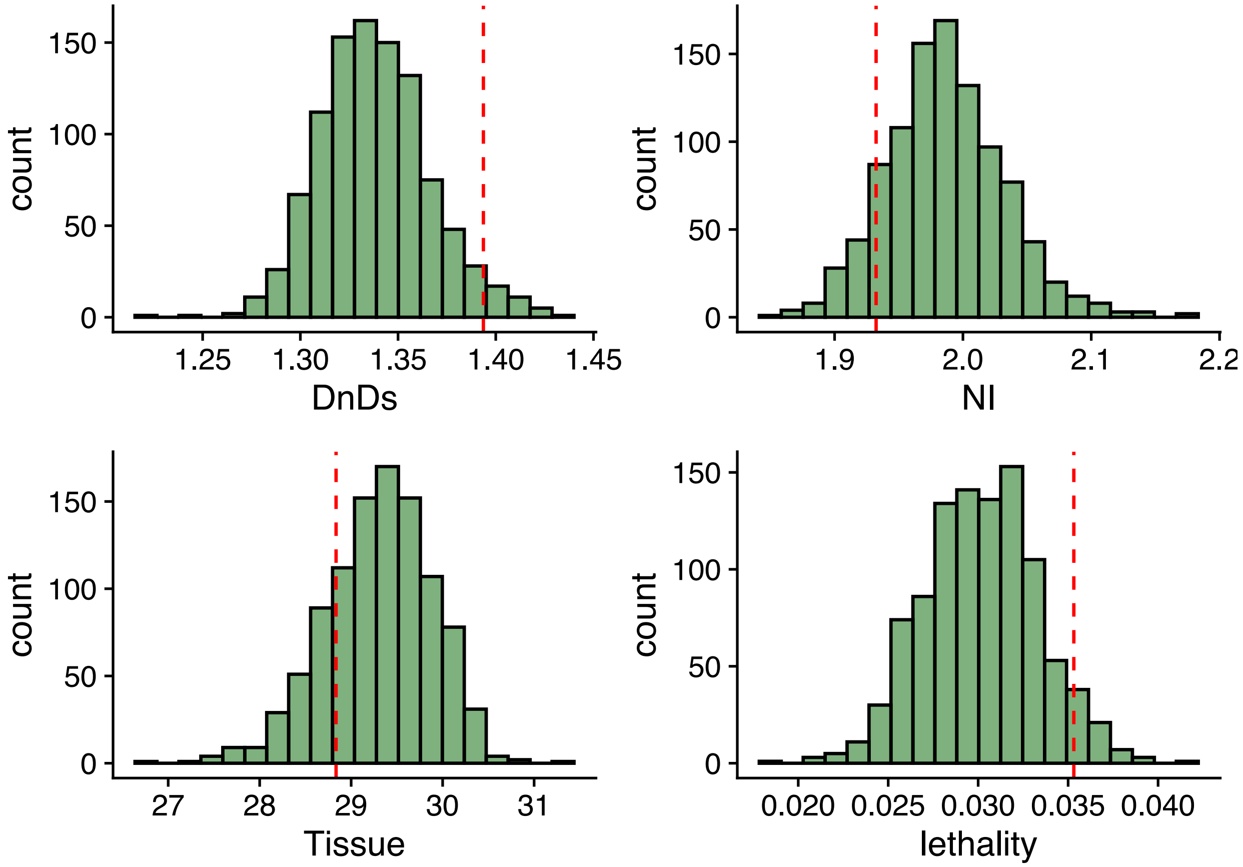

**Figure S2. Functional similarity among genes with positively covarying functional states, Molecular Function and Cellular Component.** As Figure 2B, for (A) GO Molecular Function and (B) GO Cellular Component annotations. Red dashed lines show the observed proportion of gene pairs with significant positive covariation sharing at least one term; blue distributions show 100 permutations in which gene-pair assignments were independently shuffled.

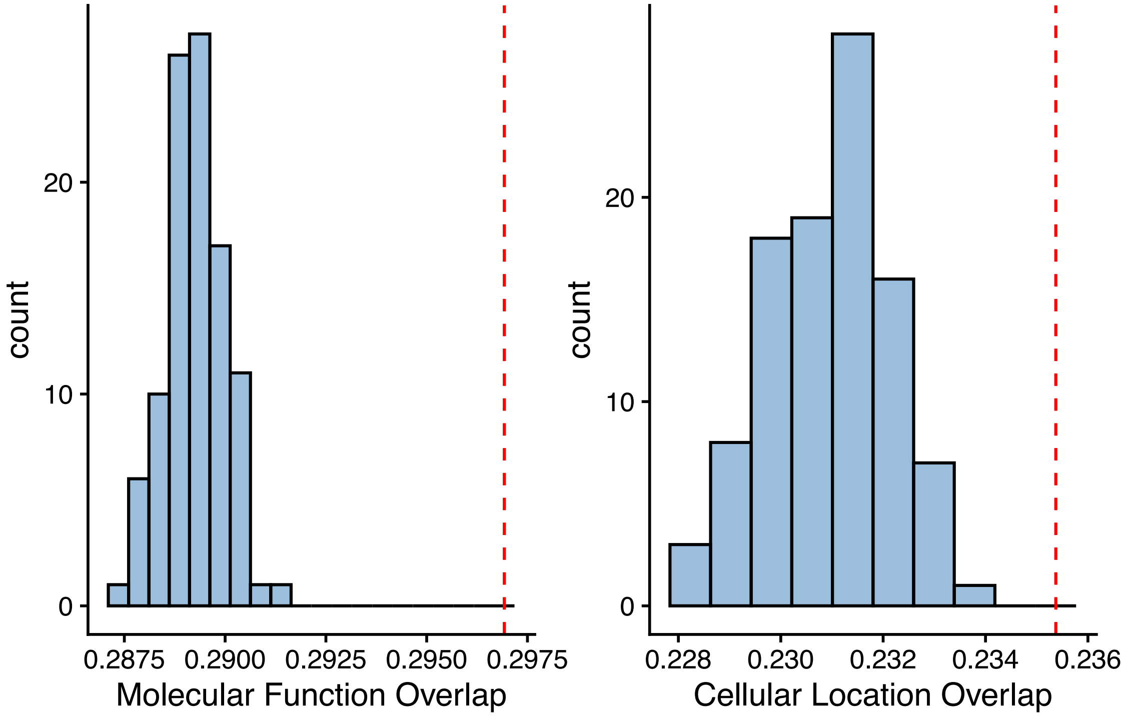

**Figure S3. Functional similarity among gene pairs showing negative covariation.** As Figures 2 and S2, computed for the 9,465 gene pairs with significant negative covariation: (A) mean coexpression, (B) GO Biological Process overlap (C) GO Molecular Function overlap, (D) GO Cellular Location overlap

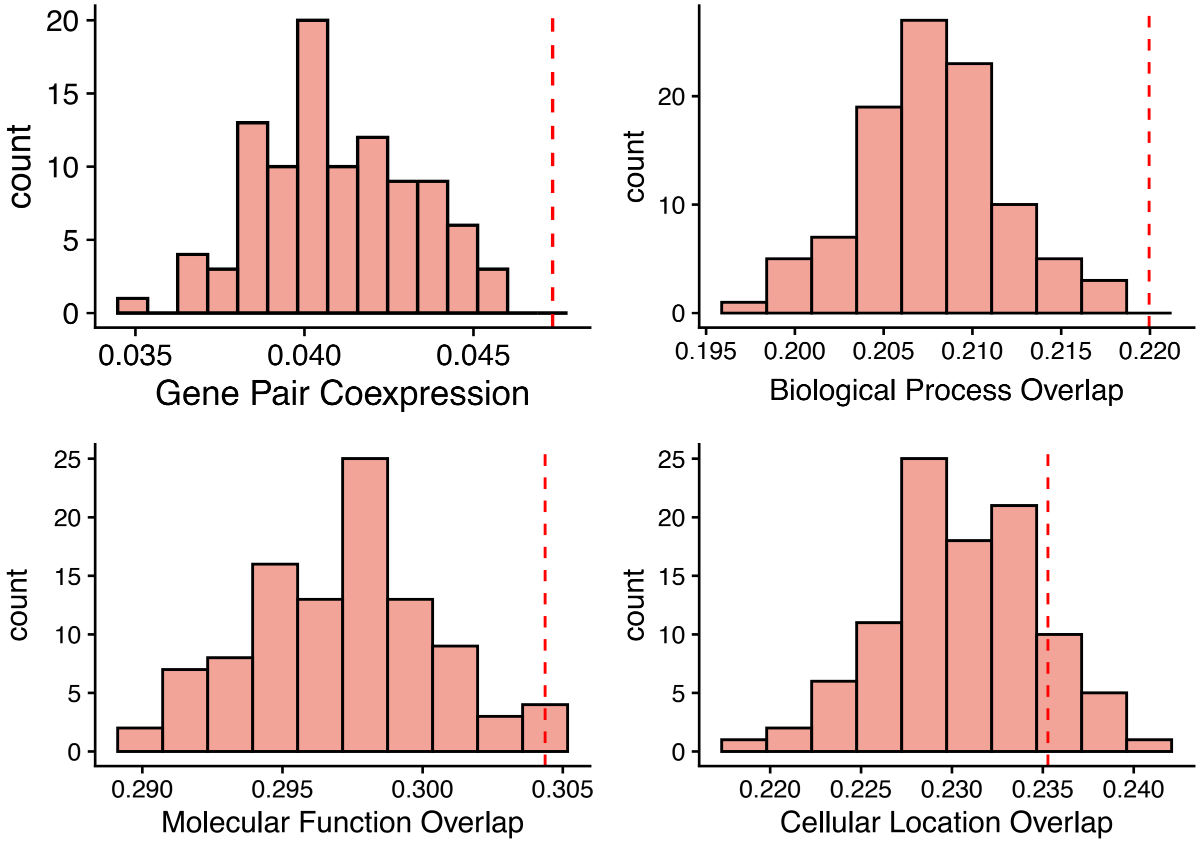

**Table S1. Accession-level associations of community loss-of-function burden.** **Panel A**, pairwise correlations between community burdens, shown as raw Pearson correlations and as partial correlations controlling for latitude and longitude. **Panel B**, correlations of each community's burden with latitude and longitude, the proportion of variance explained by admixture group (one-way ANOVA $R^2$), and correlations with accession scores on the first five principal components of the Eurasian loss-of-function matrix.

Panel A. Community-pair burden correlations

| Comparison | r (raw) | p (raw) | r (partial) | p (partial) |
|---|---|---|---|---|
| C1 vs C2 | 0.673 | 1.10E-132 | 0.684 | 2.20E-138 |
| C1 vs C3 | 0.343 | 6.60E-29 | 0.374 | 2.20E-34 |
| C1 vs C4 | 0.717 | 1.70E-158 | 0.719 | 2.70E-159 |
| C2 vs C3 | 0.397 | 5.50E-39 | 0.343 | 7.00E-29 |
| C2 vs C4 | 0.729 | 3.00E-166 | 0.727 | 1.20E-164 |
| C3 vs C4 | 0.238 | 2.40E-14 | 0.214 | 8.40E-12 |

Panel B. Community burden vs geography, admixture group and PCs

| Community | r (latitude) | p (latitude) | r (longitude) | p (longitude) | Admixture R2 | p (admixture) | PC1 | PC2 | PC3 | PC4 | PC5 |
|---|---|---|---|---|---|---|---|---|---|---|---|
| C1 | 0.055 | 0.081 | 0.059 | 0.064 | 0.382 | 4.30E-97 | 0.77 | -0.315 | 0.424 | 0.134 | 0.097 |
| C2 | 0.207 | 3.60E-11 | -0.049 | 0.12 | 0.123 | 1.40E-23 | 0.65 | -0.518 | 0.18 | -0.308 | -0.108 |
| C3 | 0.526 | 4.20E-72 | 0.006 | 0.86 | 0.489 | 2.40E-137 | 0.61 | -0.097 | -0.479 | -0.138 | 0.109 |
| C4 | 0.107 | 0.00069 | -0.011 | 0.72 | 0.069 | 8.10E-12 | 0.61 | -0.498 | 0.383 | -0.339 | -0.093 |
